# Chaperones facilitate heterologous expression of naturally evolved putative *de novo* proteins

**DOI:** 10.1101/2022.03.02.482622

**Authors:** Lars A. Eicholt, Margaux Aubel, Katrin Berk, Erich Bornberg-Bauer, Andreas Lange

**Author notes:** these authors contributed equally (Lars A. Eicholt, Margaux Aubel). corresponding author: Andreas Lange, Huefferstraße 1, 48149 Muenster, Germany.

## Abstract

Over the past decade, evidence has accumulated that new protein coding genes can emerge *de novo* from previously non-coding DNA. Most studies have focused on large scale computational predictions of *de novo* protein coding genes across a wide range of organisms. In contrast, experimental data concerning the folding and function of *de novo* proteins is scarce. This might be due to difficulties in handling *de novo* proteins *in vitro*, as most are predicted to be short and disordered. Here we propose a guideline for the effective expression of eukaryotic *de novo* proteins in *Escherichia coli*.

We used 11 sequences from *Drosophila melanogaster* and 10 from *Homo sapiens*, that are predicted *de novo* proteins from former studies, for heterologous expression. The candidate *de novo* proteins have varying secondary structure and disorder content. Using multiple combinations of purification tags, *E. coli* expression strains and chaperone systems, we were able to increase the number of solubly expressed putative *de novo* proteins from 30 % to 62 %. Our findings indicate that the best combination for expressing putative *de novo* proteins in *E.coli* is a GST-tag with T7 Express cells and co-expressed chaperones. We found that, overall, proteins with higher predicted disorder were easier to express.

## Introduction

*De novo* genes originate from intergenic or non-coding DNA regions [Tautz and Domazet-Lošo, 2011, McLysaght and Hurst, 2016, Schmitz and Bornberg-Bauer, 2017, Van Oss and Carvunis, 2019, Rödelsperger et al., 2019, Bornberg-Bauer et al., 2021, Heames et al., 2022] in contrast to genes that emerge by duplication [Liberles et al., 2011, Ohno, 1970] or rearrangement from existing gene fragments [Bornberg-Bauer and Albà, 2013]. Therefore, recent, true *de novo* genes have no precursor by definition and have not been subjected to selection for particular structures or functions for long, if at all. Due to their recent emergence, *de novo* genes tend to be shorter, evolve more rapidly and have lower expression than established genes [Van Oss and Carvunis, 2019, Schmitz and Bornberg-Bauer, 2017]. However, their short length and accelerated evolution hinder the reliable assignment of homologs. By combining homology and synteny based approaches for *de novo* gene identification, the accurate origin of *de novo* genes can be detected [Vakirlis et al., 2020].

Several *de novo* protein-coding genes have been identified and confirmed across a wide range of eukaryotes [Begun et al., 2006, Cai et al., 2008, Neme and Tautz, 2013, McLysaght and Guerzoni, 2015, Schlötterer, 2015, Schmitz et al., 2018, Vakirlis et al., 2018, Prabh and Rödelsperger, 2019, Zhang et al., 2019, Heames et al., 2020, Dowling et al., 2020]. These *de novo* genes were mainly analysed with comparative genomics and transcriptomics. A recent study by Grandchamp *et al.* (2022) showed that proto-genes, an intermediate step in *de novo* gene emergence [Domazet-Lošo et al., 2017], may already contain gene-like structures like introns, whose number and position correspond to the genomic position of proto-gene emergence. However, without experimental evidence on structure and function, our evolutionary understanding of how *de novo* proteins emerge, is incomplete.

Difficulties in handling *de novo* proteins, together with the novelty of the research area, might be the reason for the lack of experimental studies on *de novo* proteins. So far only two *de novo* proteins were expressed and characterised experimentally, Goddard (Gdrd) [Lange et al., 2021] and Bsc4 [Bungard et al., 2017]. In both cases the expressed *de novo* protein was difficult to analyse due to unstable or incorrect folding (Bsc4) or unusual behaviour in SDS-PAGE (Gdrd). Compared to well-studied proteins with expression and purification data available, *de novo* proteins tend to behave differently when using standard protocols.

Several studies, foremost some from the lab of Dan Tawfik [Tokuriki and Tawfik, 2009a,b,c, Jackson et al., 2022], inspired us to apply co-expression with chaperones to achieve soluble expression of *de novo* proteins. Since *de novo* proteins evolve rapidly by becoming coding from scratch, they probably lack a stable structural configuration and contain high amounts of disorder [Van Oss and Carvunis, 2019, Schmitz and Bornberg-Bauer, 2017]. Those properties determine the levels of soluble and insoluble fractions of a protein during *in vitro* experiments and could explain the obstacles faced during their expression [Soskine and Tawfik, 2010, Tretyachenko et al., 2017]. On the other hand, it is not yet clear if *de novo* proteins undergo a similar hindrance in their native organism or only in the expression hosts [Gasser et al., 2008]. While Tawfik and colleagues used chaperones to explore the sequence space of enzymes and enable soluble expression of mutants [Tokuriki and Tawfik, 2009a,b,c], we hypothesised that *de novo* protein expression might also profit from chaperones. With their “emergence from dark genomic matter” in the DNA [Bornberg-Bauer et al., 2015] and predicted lack of stability and high disorder, *de novo* proteins are prospective targets for for chaperones because their solubility can be increased. [Tokuriki and Tawfik, 2009a,b]. Increased solublity can be relevant for protein purification and any follow-up experiments.

The chaperonin GroEL and its co-chaperone GroES are found throughout the bacterial domain, while their homologs, HSP60 and HSP10, respectively, are found in eukaryotes [Finka et al., 2016]. GroEL/ GroES play a pivotal role in the translocation, dis-aggregation, function and folding of newly synthesised peptides after translation [Tokuriki and Tawfik, 2009a, Finka et al., 2016, Libich et al., 2015, Lin et al., 2008].

The other chaperone system used here is DnaK, DnaJ and GrpE (homologous to HSP70 and HSP40 in eukaryotes). For simplicity we will refer to the chaperone system GroEL/ GroES as only GroEL and to DnaK, DnaJ and GrpE as DnaK only. While the GroEL system targets misfolded and unfolded proteins, DnaK can refold an already aggregated protein to its native state using ATP (see **Figure 1**) [Schröder et al., 1993, Sharma et al., 2010, Kim et al., 2013, Mashaghi et al., 2016]. The two different chaperone systems can be exploited for challenging heterologous expression of proteins which are foreign to the host, and thus prevent misfolding and aggregation which is often associated with heterologous expression [Goloubinoff et al., 1989, Finka et al., 2016, Kim et al., 2013, Tokuriki and Tawfik, 2009a,b,c]. For this study, we used **21** putative *de novo* proteins, 11 from *Drosophila melanogaster* (termed here as *DM*1-10 and Atlas) and 10 from *Homo sapiens* (termed here as *HS*1-10) as shown in **Figure 2**. These *de novo* proteins have been recently published by Heames *et al.* and Dowling *et al.* Additionally, we tested our method on a recently published and better characterised putative *de novo* protein from *D. melanogaster*, called Atlas. Atlas appears to function as a DNA binding protein that facilitates the packaging of chromatin in developing *D. melanogaster* sperm [Rivard et al., 2021]. Since experimental work with *de novo* proteins is still underrepresented (compared to computational studies) and challenging, we want to propose a guideline for successful expression of putative *de novo* proteins in *E. coli*. We combined different chaperone systems (GroEL and DnaK) with different combinations of *E.coli* strains (BL21 Star™ (DE3) and T7 Express) in order to express putative *de novo* proteins solubly. To verify successful expression of target proteins, western blots were performed and samples sent for tryptic digest followed by mass spectrometry. We identified the best combination for expression of putative *de novo* proteins in *E. coli* and increased the total number of solubly expressed putative *de novo* proteins to 62 % (13/21 proteins). The different chaperone systems increased or enabled soluble expression in four cases (31 %), while DnaK only helped in two, GroEL in all of those four.

**Figure 1:**
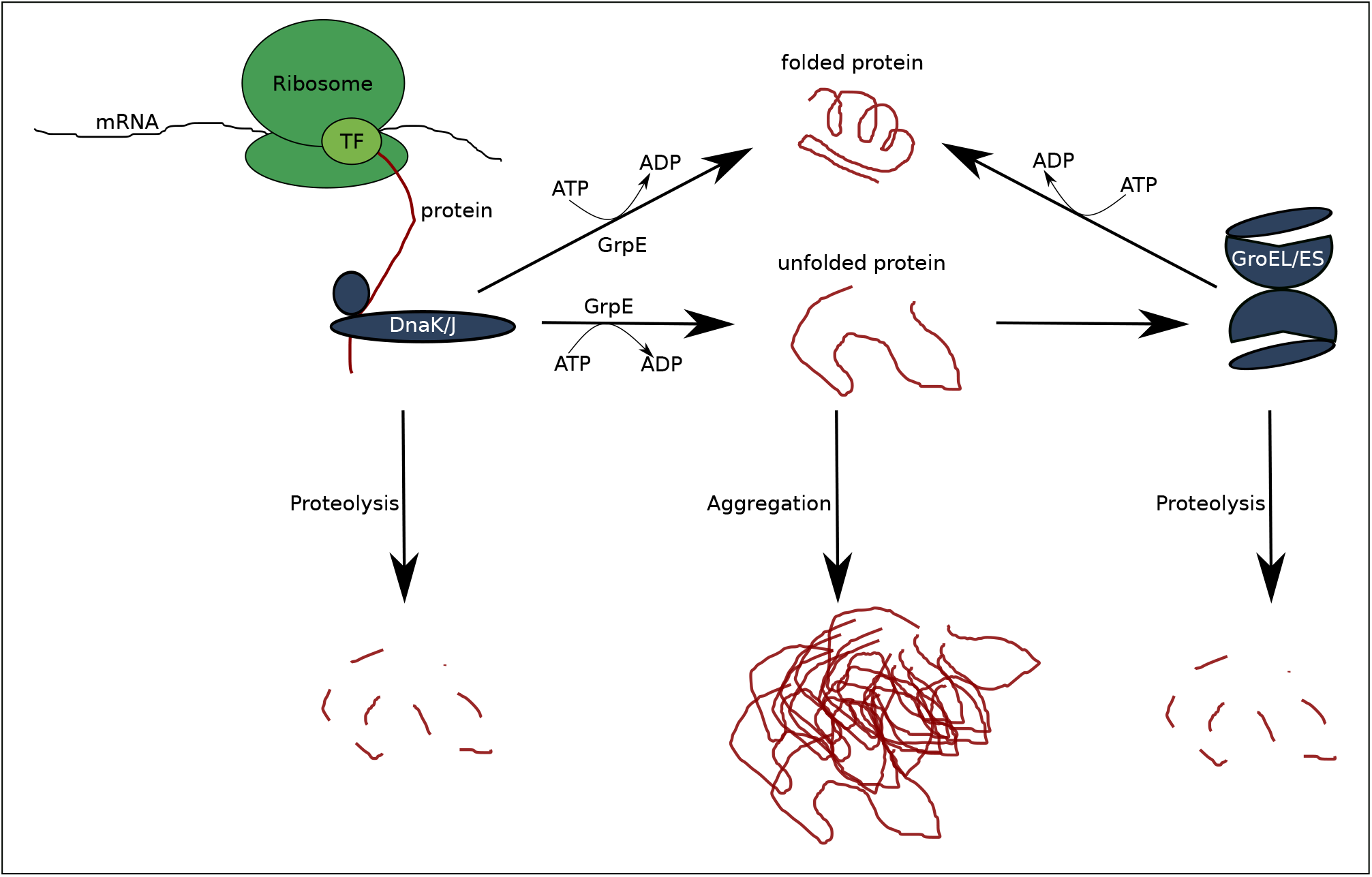
Mechanism of chaperone assisted protein folding after Thomas et al. [Thomas et al., 1997]. The nascent protein is bound by the DnaK/J complex and release is triggered by GrpE under ATP hydrolysis. After release, the protein is either correctly folded, degraded (proteolysis) or remains unfolded. The unfolded protein can either aggregate or bind to the GroEL/ES complex. GroEL/ES either releases the folded protein by ATP hydrolysis or the protein is degraded.

**Figure 2:**
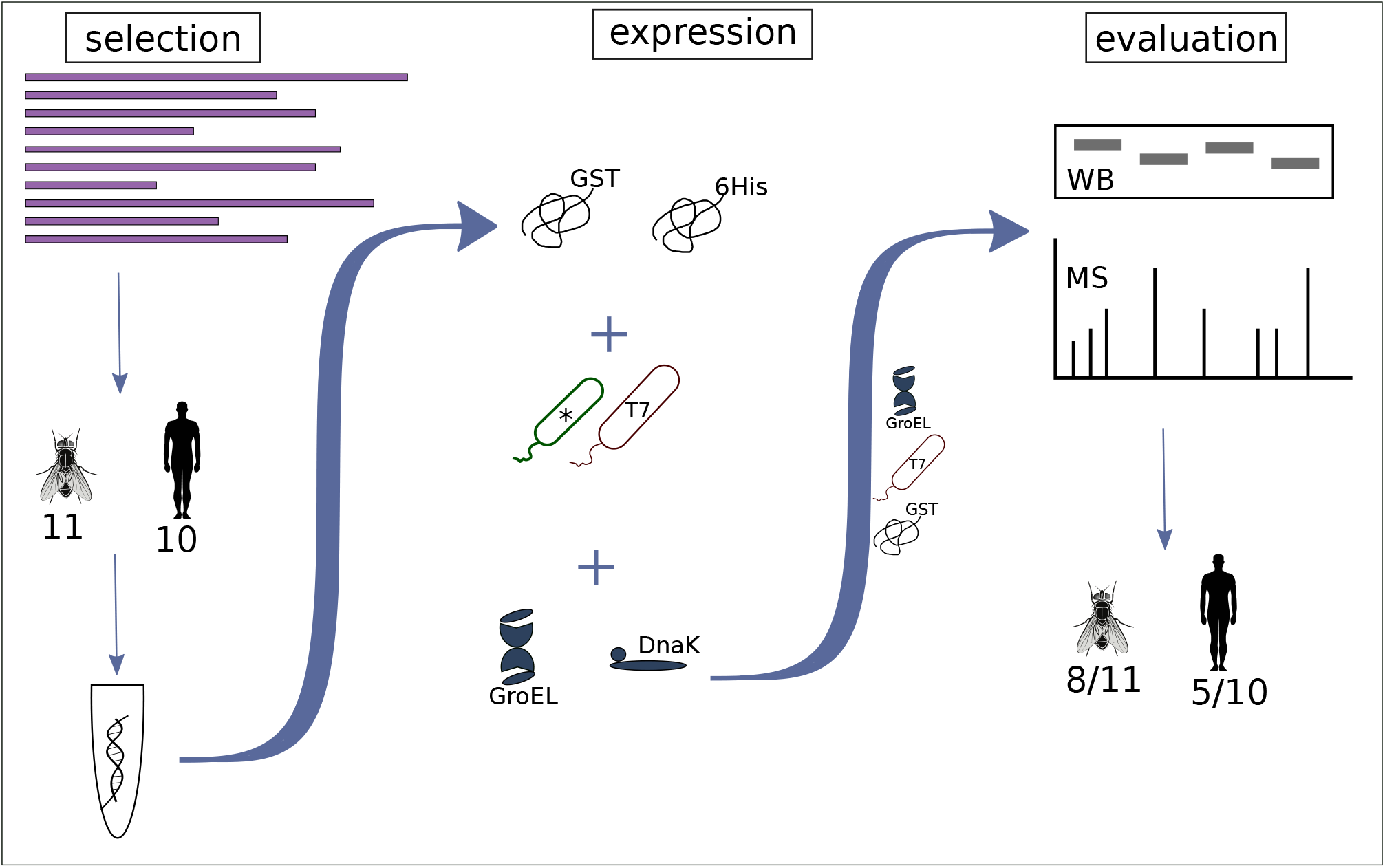
Overview of the workflow on *de novo* protein expression: We first selected candidate proteins from *Drosophila melanogaster* (11, including Atlas) and 10 from *Homo sapiens* from a pool of putative *de novo* genes for expression. The 21 sequences were codon optimised for *E. coli* and ordered from Twist. For expression, different tags (GST and His), different *E. coli* expression cells (star™, T7) and different chaperones (GroEL and DnaK systems) were tested. The success of protein expression was verified by western blot (WB) and mass spectrometry (MS). With the help of GST-tag and chaperone system GroEL using specialised T7 express cells, we could express around 50 % of the candidate *de novo* proteins solubly.

## Results

### Structural content of the putative *de novo* proteins

#### Disorder Predictions

We performed disorder predictions with IUPred2a [Erdős and Dosztányi, 2020, Mészáros et al., 2018] on all candidate *de novo* proteins. For this we calculated the percentage of residues predicted to be disordered (**Figure 3**), as opposed to the overall average disorder score (**Figure S1**). This allows direct comparison to secondary structure predictions (**Figure 4**). Our first objective here was to choose candidate *de novo* proteins with different levels of intrinsic disorder to observe any difference in their ability to express. If any trend in predicted disorder and soluble expression or susceptibility to chaperones was observed, this could help choosing promising candidates for characterisation in future experiments. The predicted disorder ranged from around 3 % to 100 % as shown in **Figure 3**. *DM*5 was predicted to have least disorder content, while *DM*6, *DM*3, *HS*10 and *DM*8 appear to be entirely disordered. The putative *de novo* protein Atlas has predicted disorder of around 60 %.

**Figure 3:**
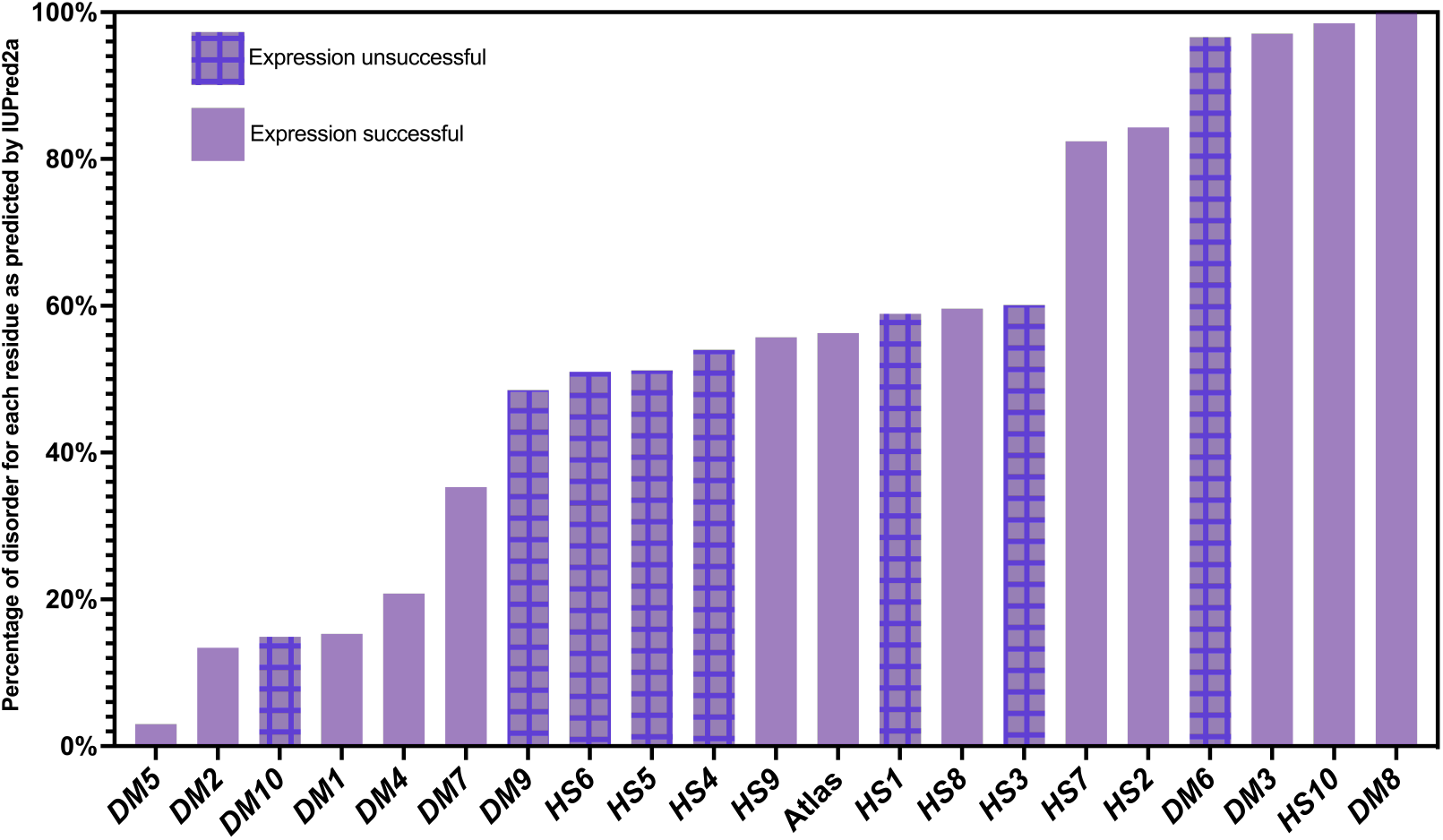
Percentage of disorder as calculated with IUPred2a. All candidate *de novo* proteins used for expression experiments ordered by their disorder level from left to right. Unicolor bars belong to the successfully expressed proteins, checked bars to the unsuccessful ones.

**Figure 4:**
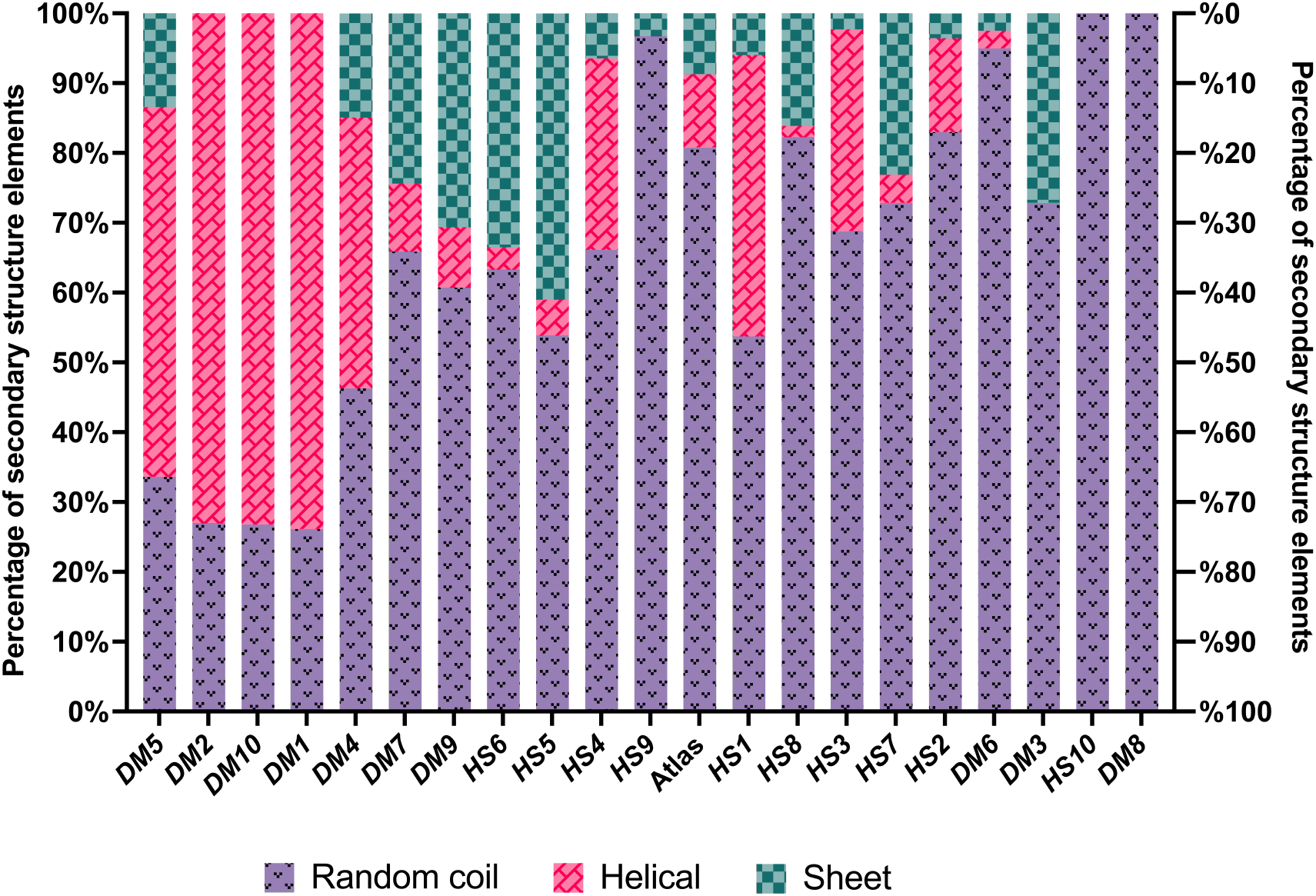
Percentage of random coils, α-helices and β-sheets predicted by Porter 5.0 for each *de novo* protein candidate. Left to right following increasing disorder level based on **Figure 3**.

#### Secondary Structure Predictions

Predictions of secondary structure elements were performed using Porter 5.0 [Torrisi et al., 2018, 2019] for all candidate proteins and are shown in **Figure 4**. While the results indicate a high amount of random coils for most candidates, they do not completely follow the trend of the disorder predictions by IUPred2a (compare **Figure 3**). *DM*3 for example, is predicted to be ~ 100 % disordered by IUPred2a, while its on the other hand predicted to have over 20 % β-sheet and ~ 70 % random coils by Porter 5.0. Our goal was to choose a cohort of *de novo* proteins that consist of a diverse range in composition of structural elements.We assumed that a protein containing more secondary structure elements should be better accessible for soluble expression with chaperones. Notably, *DM*1, *DM*2, *DM*4, *DM*5 and *DM*10 are predicted to have secondary structure contents of 50 % or more, with α-helices to be more frequent than β-sheets. *HS*4, *HS*5, *HS*6; *HS*7, *DM*3 and *DM*7, on the other hand, are predicted to be mostly random coils (disordered) with otherwise high amounts of β-sheets predicted.

### Expression of putative *de novo* proteins

#### Candidates of *Drosophila melanogaster*

Our initial approach was similar to the successful expression of characterised putative *de novo* protein Gdrd [Lange et al., 2021]. Therefore, we aimed to express our **11** putative *de novo* protein candidates with an N-terminal 6x His-tag in *E. coli* BL21 Star™ (DE3) cells, and verify expression via SDS-PAGE and mass spectrometry. However, for our candidates the expression level was either very low or not detectable, as can be seen in **Figure S3**. We switched to different *E. coli* cells (T7 Express), but expression remained unsuccessful. Shifting from an N-terminal 6xHis-Tag to a C-terminal 6xHis-tag showed similar negative results. Considering the size and levels of disorder, we switched to a larger tag for increased solubility and stability, choosing an N-terminal GST-tag. In this way we were able to observe a higher success rate in soluble expression of our target proteins. But not all proteins could be expressed at satisfying levels, especially solubility needed to be increased for some (**Figure S3**).

Inspired by successful work carried out by Tawfik *et al.* (2009a, 2009b, 2009c) we hypothesised that chaperones could improve thermodynamic stability of these evolutionarily young proteins thus enabling their soluble expression. We repeated our experiments with the addition of the two chaperone systems i) GroEL and ii) DnaK. We were able to increase the number of solubly expressed *de novo* candidate proteins of *D. melanogaster* using the combination of either GroEL or DnaK and N-terminal GST-tag (see **Figure 5**). However, for the candidate proteins *DM*6, *DM*9, and *DM*10 no soluble expression was achievable, despite the use of different tags, strains, or chaperones. Only in the case of Atlas, the combination of N-terminal 6x His-tag and GroEL worked best. We tested all combinations in BL21 Star™ (DE3) and T7 Express *E.coli* cells. Six candidate proteins were expressed in T7, two were expressed in BL21 Star™ (DE3) cells. Three proteins were not expressable in either strain. In summary, with the combination of chaperones and switching to N-terminal GST-tag, we were able to express 73 % of the *D. melanogaster* putative *de novo* protein candidates (see **Table 1**).

**Table 1:**
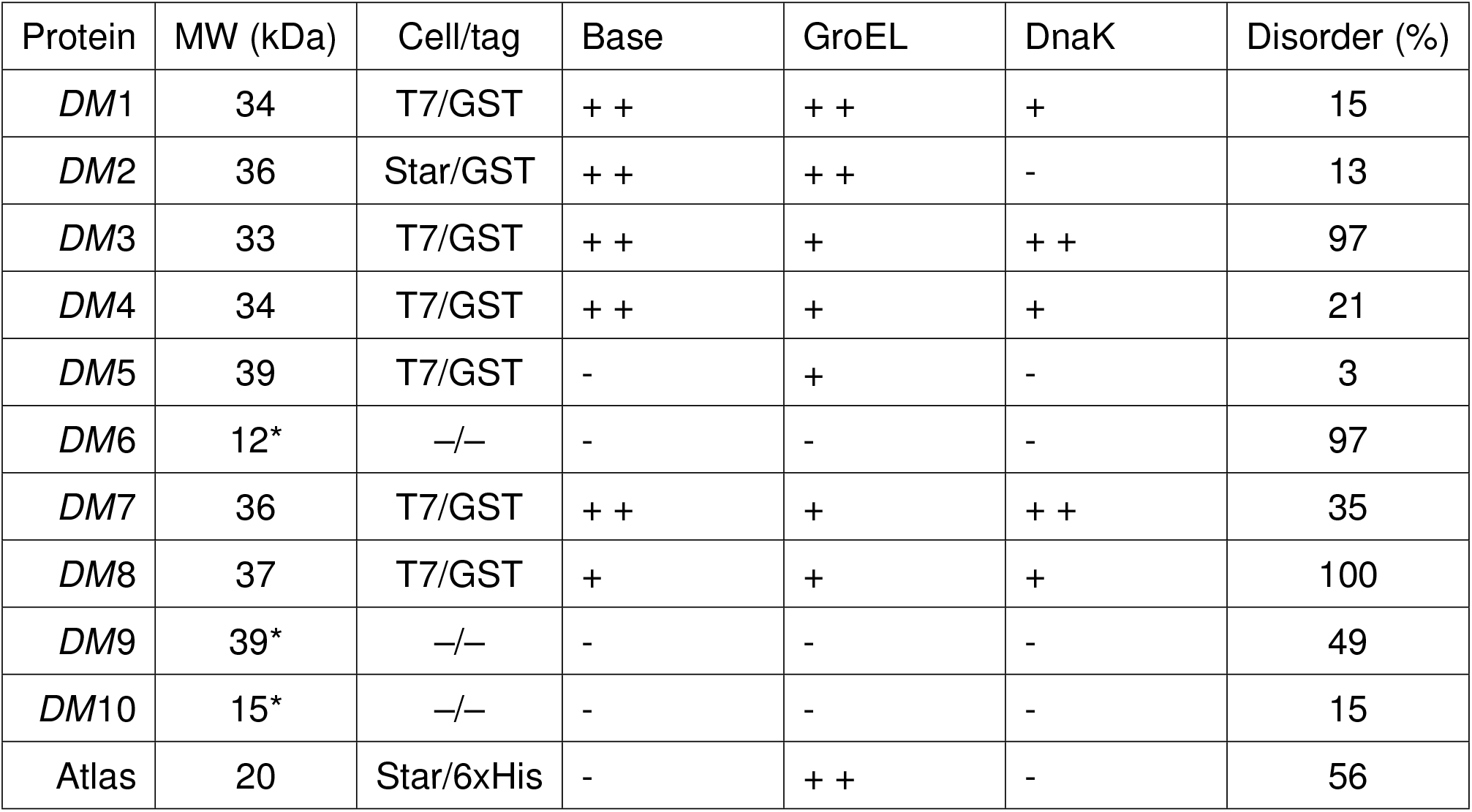
Expression conditions & results of *D. melanogaster de novo* proteins. Base = no chaperones, GroEL = GroEL/ES, DnaK = DnaK/J/GrpE. *Molecular weight (MW) without tag. Plus signs mean visible expression, two plus signs strong expression, minus sign means no visible expression.

**Figure 5:**
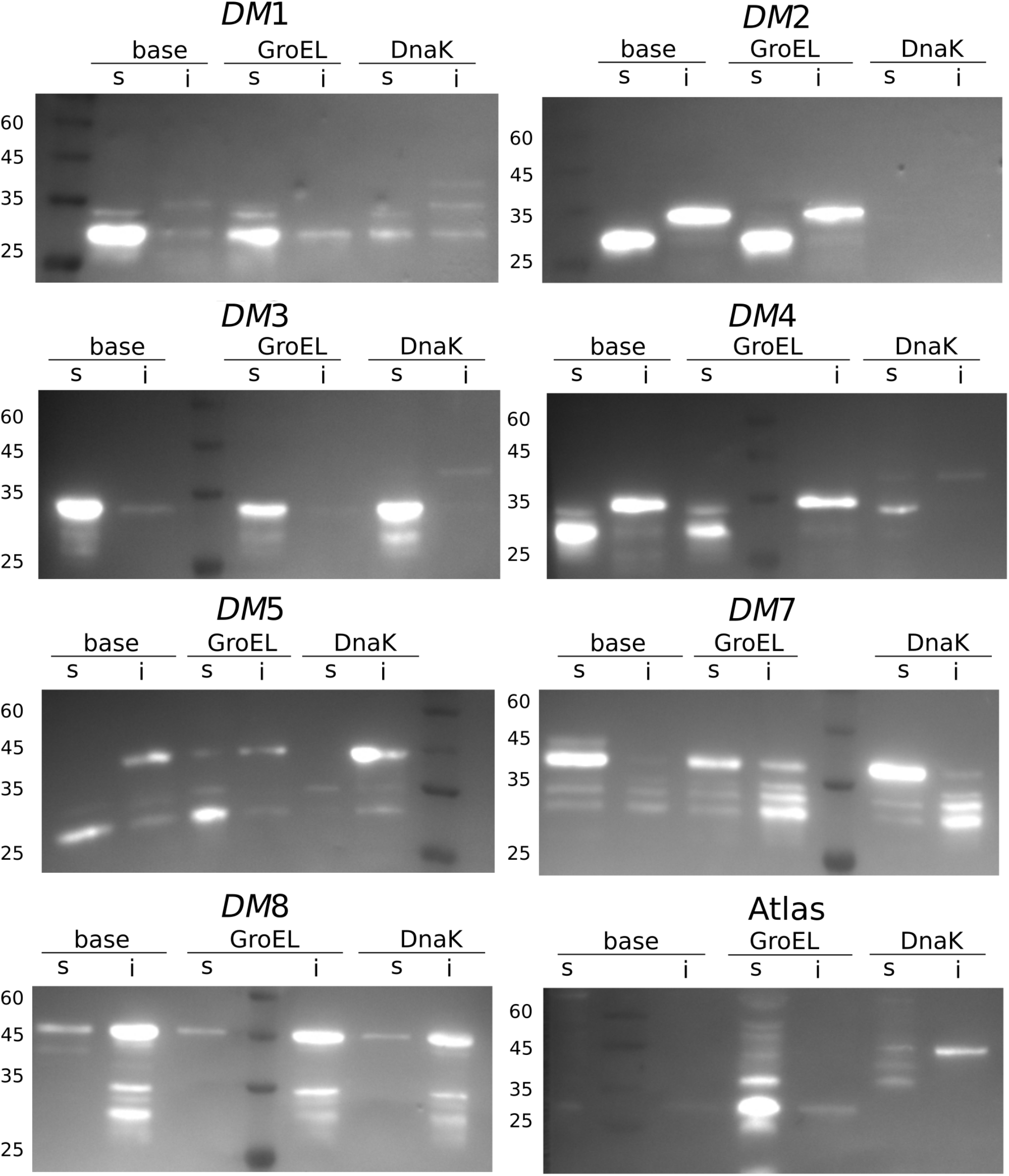
Western blots with Anti-His antibody: ***DM*1** (34 kDa): highest solubilty without chaperones, then GroEL, then DnaK; highly soluble. ***DM*2** (36 kDa): only insoluble, even with chaperones. ***DM*3** (33 kDa): DnaK highest solubilty, then base, then GroEL; very soluble. ***DM*4** (34 kDa): DnaK highest solubilty, then GroEl, then base; very insoluble. ***DM*6** (39 kDa): GroEL only one with soluble fraction, runs a bit high. ***DM*7** (36 kDa): Dnak highest solubilty, then base, then GroEL very soluble. ***DM*8** (37 kDa): all similar, different expression levels, first base, then GroEL, then DnaK; more insoluble. **Atlas** (20 kDa): GroEL highest solubilty, nothing in base and DnaK.

#### Comparison of different chaperone conditions for *D. melanogaster* proteins

Western blots were used for comparison of the soluble expression levels with and without chaperones, in order to test our hypothesis that chaperones would increase soluble expression of the target proteins. The optimal conditions identified by SDS-PAGEs were repeated under three settings: i) without chaperones (base), ii) with GroEL and iii) with DnaK. Surprisingly, we did not observe increased solubility for most putative *de novo* proteins when adding chaperones (see **Figure 5** and **Table 1**).

In contrast, we observed soluble expression for most proteins without chaperones, e.g. *DM*1, *DM*2, *DM*3, *DM*4 and *DM*7. In combination with GroEL, the intensity of the bands in the soluble fraction, and therefore amount of soluble protein, even decreased for *DM*3, *DM*4 and *DM*7. For *DM*2 and *DM*5 the amount of soluble protein increased when coexpressed with GroEL. When DnaK was co-expressed, protein solubility either appeared to decrease (*DM*1, *DM*2 and *DM*4), or was similar to the base (*DM*3 and *DM*7). *DM*8 showed similar soluble expression for all three conditions with most of the protein being insoluble. In the case of Atlas and *DM*5, soluble protein expression was increased or enabled with the addition of the GroEL chaperone system while DnaK and base expression resulted in no or very little soluble protein. While we cannot confirm that co-expression with DnaK in fact decreases the amount of soluble protein (*DM*1, *DM*2 and *DM*4), we do not see increased soluble expression for any of the candidate proteins in the presence of DnaK as we do for GroEL (*DM*5 and Atlas).

#### Candidates of *Homo sapiens*

The 10 putative human *de novo* proteins were expressed following the same protocol as the *D. melanogaster* proteins by combining the different *E. coli* expression cells, tags and chaperone systems (**Figure S4**). We detected a similar trend here as for the *D. melanogaster* proteins (N-terminal GST-tag in *E. coli* T7 express cells; see **Table 2**). One protein (*HS*8), however, was only weakly expressed with an N-terminal 6xHis-tag but using also *E. coli* T7 express cells. Without the addition of chaperones only *HS*7, *HS*8 and *HS*10 were successfully expressed and soluble. After co-expression with chaperones, as described for *D. melanogaster* proteins, two more *H. sapiens* proteins could be expressed. Unfortunately, *H. sapiens* protein candidates *HS*1, *HS*3, *HS*4, *HS*5 and *HS*6 showed no expression at all, even with chaperones. In total we were able to express 5 out of 10 putative *de novo* proteins following our protocol (see **Table 2**).

**Table 2:** Expression conditions & results of *H. sapiens de novo* proteins. Base = no chaperones, GroEL = GroEL/ES, DnaK = DnaK/J/GrpE. *Molecular weight (MW) without tag. Plus signs mean visible expression, two plus signs strong expression, minus sign means no visible expression.

| Protein | MW (kDa) | Cell/tag | Base | GroEL | DnaK | Disorder (%) |
| --- | --- | --- | --- | --- | --- | --- |
| *HS1* | 17* | —/— | - | - | - | 59 |
| *HS2* | 44 | T7/GST | - | + | + | 84 |
| *HS3* | 20* | —/— | - | - | - | 60 |
| *HS4* | 16* | —/— | - | - | - | 54 |
| *HS5* | 15* | —/— | - | - | - | 51 |
| *HS6* | 22* | —/— | - | - | - | 51 |
| *HS7* | 50 | T7/GST | + + | + + | - | 82 |
| *HS8* | 16 | T7/6xHis | + | + | + | 60 |
| *HS9* | 42 | T7/GST | - | + | + | 56 |
| *HS10* | 43 | T7/GST | + | + | + | 99 |

#### Comparison of different chaperone conditions for *H. sapiens* proteins

Western blots were used for comparison of the three different chaperone expressions i) base, ii) GroEL and iii) DnaK, as described above. Two out of the five successful candidates (*HS*2 and *HS*9) showed very weak or no soluble expression without chaperones, but solubility could be increased with both chaperone systems. *HS*8 and *HS*10 showed low soluble expression overall, but no change in solubility was visible when co-expressing with either chaperone system. The candidate *de novo* protein *HS*7 already showed strong soluble expression at base (**Figure 6**). However, the addition of GroEL seemed to increase soluble expression further, while DnaK co-expression led to low or no protein being detected. Overall, the trend observed for the *D. melanogaster* proteinś was consistent with the trend observed for the *H. sapiens* proteins. GroEL increased soluble expression for most putative *de novo* proteins while DnaK lacked substantial influence on protein solubility.

**Figure 6:**
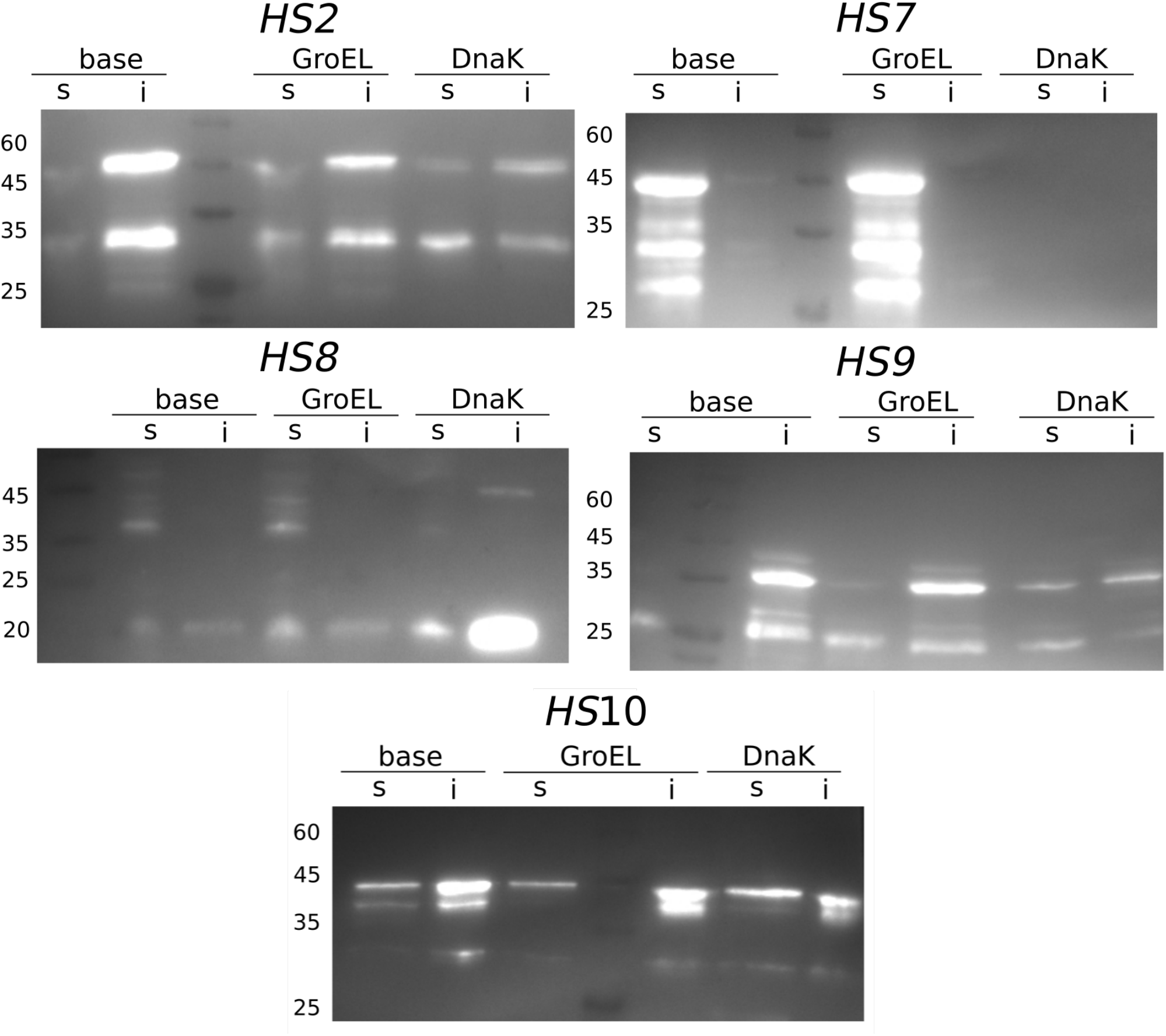
Western blots with Anti-His antibody: ***HS*2** (44 kDa): upper bands (lower are degraded protein or double bands) most in DnaK, then GroEL, then base; very insoluble. ***HS*7** (50 kDa): GroEL best, then base, nothing in DnaK. Possible protein degradation; very soluble. ***HS*8** (16 kDa): upper bands most in DnaK, then GroEL, then base; very insoluble. ***HS*9** (42 kDa): upper bands (lower are degraded protein or double bands) most in DnaK, then GroEL, then base; very insoluble. ***HS*10** (43 kDa): upper bands (lower are degraded protein or double bands) most in GroEL, then DnaK, then base; very insoluble.

## Discussion

*De novo* proteins have first been detected more than a decade ago and the mechanism of their emergence has been studied intensely ever since [Begun et al., 2006, Van Oss and Carvunis, 2019]. Still, there are concerns (i) regarding the reliability of their computational identification [Moyers and Zhang, 2015, Domazet-Lošo et al., 2017, Weisman et al., 2022] and (ii) if and how they code for functional proteins. To shed light on these concerns, *de novo* proteins need to be studied experimentally as well as theoretically. The handling of *de novo* proteins by heterologous expression and purification is often difficult because solubility is low and purification yields little amounts and potentially unstable proteins. Moreover, identifying the function of these young genes, is another challenging task. In this study we present a guideline for expressing *de novo* proteins in *E. coli*.

### Expression cells

*E. coli* is the most widely used model organism for recombinant expression. However, foreign proteins can be toxic to *E. coli* by interfering with the physiology or leading to protein aggregation. This may result in low expression yields, growth defects or even cell death (Saïda et al. [2006], Saïda [2007], Rosano and Ceccarelli [2014]). Optimized expression hosts and plasmids (Saïda et al. [2006], Saïda [2007], Rosano and Ceccarelli [2014]) or chaperones can be used to overcome the expression issues caused by proteins which are a metabolic burden for the host. Here, we used two different types of the *E. coli* strains (DE3): BL21 Star™ and T7 Express. Both strains resulted in effective protein expression and a relatively high yield of the *de novo* proteins, with T7 Express being the best option. The *de novo* proteins studied here are possibly a toxic, metabolic burden to the *E. coli* cells, suggesting T7 cells are the better choice of expression cell. BL21 Star™ (DE3) contains a T7-RNA-polymerase under control of lacUV5 promoter together with higher mRNA stability. This leads to stable mRNA transcripts and higher amount of target protein. However, BL21 Star™ (DE3) cells have increased basal expression of heterologous genes and cannot express toxic genes. In contrast, the T7 Express cells have a reduced basal expression of target proteins than BL21 Star™ (DE3) cells. Therefore, toxic proteins can be expressed better in T7 cells compared to BL21 Star™ [New England Biolabs].

### Comparing different protein tags

Based on our study, an N-terminal GST-tag was the more appropriate choice than a 6x Histag. Some *de novo* protein candidates are quite small (8-12 kDa), so a larger tag like GST might already stabilise in a chaperone-like manner [Harper and Speicher, 2011, Rosano and Ceccarelli, 2014]. However, Atlas and *HS*8, i.e. two out of 21, were only expressed with an N-terminal 6x His-tag. With a mass of only 1 kDa, 6x His-tag is the better choice for further structural characterisation using circular dichroism (CD), multi-angle light scattering (MALS) or nuclear magnetic resonance (NMR), since a small tag has less influence on protein folding. In contrast, the larger GST-tag needs to be cleaved for most follow-up experiments. When removing the tag, the *de novo* protein might behave differently and could degrade or aggregate.

### Influence of chaperones on protein expression and solubility

Our western blot results indicate that GroEL slightly outperforms DnaK in terms of increased protein solubility. In some cases both chaperone systems increase or enable soluble expression (*HS*2 and *HS*9, 2/21) but for most proteins GroEL leads to more soluble protein than DnaK (*DM*1, *DM*2, *DM*5, Atlas and *HS*7, 5/21). DnaK requires easily accessible hydrophobic fragments that can be predicted from the protein sequence, while GroEL demands no defined binding motifs. However, in the case of our proteins we found no connection between predicted DnaK binding sites and influence of DnaK on protein expression level (**Figure S2**). Contrary to our findings here, we did not observe that GroEL increases protein solubility in Heames et al. [2022], where we used a library of 1800 putative *de novo* proteins (4 - 8 kDa) in a cell-free expression system.

We cannot verify that changes with co-expression of chaperones is solely due to effects of chaperones on putative *de novo* proteins or on overall amount of protein expression. Our main interest here is to optimise expression for follow-up experiments and not to draw general conclusions on chaperone interaction with *de novo* proteins. Drawing conclusions from heterologous expression experiments towards *in vivo* interactions of proteins and chaperone systems is fragmentary and can only serve as hypotheses in need of further verification using *in vivo* experiments [Niwa et al., 2012].

### Comparing putative *de novo* proteins from *D. melanogaster* to *H. sapiens*

In total we were able to successfully express 13 out of 21 putative *de novo* proteins in *E. coli* cells (eight in *D. melanogaster* and five in *H. sapiens*), resulting in a success rate of 62 %. For both, *D. melanogaster* and *H. sapiens* candidate putative *de novo* proteins, the combination of GST-tag and *E. coli* T7 Express cells were the best performing (10 out of 13; 77 %). We performed test expressions and compared the levels of soluble expression for different chaperone combinations shown in **Figures 5** and **6**. Expression results from putative *de novo* protein candidates *DM*5, Atlas, *HS*2 and *HS*9 were in line with our original hypothesis that chaperones enhance solubility of *de novo* proteins in heterologous expression systems. However, the choice of appropriate tag and expression cells in the first step was equally, if not more, important. When using the N-terminal His-tag that proved successful for putative *de novo* protein Gdrd, only two (Atlas and *HS*7) of our candidate proteins were expressed. When switching to the N-terminal GST-tag another seven *D. melanogaster* and four more *H. sapiens* protein candidates were expressed. Unfortunately, we were not able to express 38 % of the candidate proteins in *E. coli* at all (*HS*1, *HS*3 - *HS*6, *DM*6, *DM*9 and *DM*10), despite trying different expression strains, tags and chaperone systems.

### Disorder & secondary structure predictions

When examining the predicted structural properties of the human *de novo* protein candidates, we observe a slight trend towards better expression of the more disordered proteins. This trend can be observed for the IUPred2a disorder predictions (**Figure 3**) but becomes more apparent for the overall secondary structure predictions (**Figure 4**). The unsuccessful expression candidates *HS*1, *HS*3 and *HS*4 showed a higher predicted α-helical content of approximately 40 % while *HS*5 and *HS*6 had a higher predicted β-sheet content of around 30 - 40 % compared to the other human candidate proteins. The described differences in predicted secondary structure content and disorder level might be the reason why these putative *de novo* candidates could not be expressed in *E. coli* cells even with the help of chaperones.

For *D. melanogaster* protein candidates, this trend was not observed. Here, several of the proteins with lower disorder predicted (*DM*1, *DM*4 and *DM*7) were expressed solubly without addition of chaperones. Yet, *DM*6 (~ 90 % disorder predicted) was not expressed successfully. However, the two proteins with 100 % random coils predicted by Porter 5.0 and highest disorder predictions by IUPred2a (*DM*8 and *HS*10) did not show any change in solubility when chaperones were co-expressed. Considering that such highly disordered proteins do not need chaperones, this observation was expected.

Deviations of the level of predicted disorder and predicted secondary structures, especially random coils, for each protein can be explained by the differences of IUPred2a and Porter 5.0. IUPred2a provides energy estimations for each amino acid residue resulting in quasi-probabilities of disorder [Mészáros et al., 2018]. On the other hand, Porter 5.0 is based on a neural network relying on sequence alignments and co-evolutionary information [Torrisi et al., 2018]. These fundamentally different approaches can lead to inconsistent results in some cases (e.g. *HS*9, *DM*3) while not invalidating one another.

## Conclusion and Outlook

Exemplifying the general trend for soluble *de novo* protein expression is only the first step towards enabling further *in vitro* experiments for functional and structural characterisation. Further advancement will lead to efficient and stable purification, followed by functional assays such as peptide phage display to identify binding partners [Sundell and Ivarsson, 2014, Ivarsson et al., 2014]. This technique has proven to be useful for high-throughput screening of intrinsically disordered regions for short linear motifs [Ali et al., 2020], especially for human proteins. Soluble expression and purification will be crucial for structural characterisation via CD, NMR and Cryo-EM. Due to their small size and high disorder content, only NMR [Lange et al., 2021] and potentially Cryo-EM [Chiu et al., 2021] will be capable of solving the structure of *de novo* proteins experimentally. Even in light of the recent dawn of computational structure prediction [Jumper et al., 2021, Baek et al., 2021], experimental structural and functional determination remains necessary, especially for *de novo* proteins. While contemporary prediction methods can certainly provide a first estimate on structure, the intrinsic nature of *de novo* proteins, with their short length, high disorder content and lack of homology, will demand some scepticism while analysing such predictions [Ruff and Pappu, 2021, Monzon et al., 2022].

This study of 21 putative *de novo* proteins from *H. sapiens* and *D. melanogaster*, including previously *in vivo* characterised putative *de novo* protein Atlas, showed that chaperones may help expressing *de novo* proteins in *E. coli* cells. However, not all putative *de novo* proteins needed chaperones for soluble expression and sometimes even expressed better without. Fusion of the target *de novo* proteins to a GST-tag and using T7 Express cells as hosts proved to be the most successful combination. Our work may serve as a guide to facilitating future analyses of putative *de novo* proteins or other difficult (short and/or disordered) target proteins in *E. coli*.

## Material and Methods

### Online data availability

All SDS-PAGEs, MS results, western blots and scripts are deposited in Zenodo database (https://doi.org/10.5281/zenodo.6283284)

### Computational Methods

#### Candidate selection

We selected a total of 21 putative *de novo* protein candidates. Ten are uncharacterised putative *de novo* proteins from *Homo sapiens* [Dowling et al., 2020] and are referred to here as *HS*1-10. Ten proteins originate from *Drosophila melanogaster* [Heames et al., 2020] and are referred to as *DM*1-10. One is the functionally characterised putative *de novo* protein Atlas from *D. melanogaster* [Rivard et al., 2021]. The 21 candidates contain different levels of disorder and secondary structure elements (α-helix, β-sheet, mixture of both) and different sequence lengths (see **Figure 3**). We selected only candidate sequences without exon/intron structure and without long single amino acid repeats. All putative *de novo* proteins have confirmed expression in their native organism.

#### Predictions

We performed disorder predictions with IUPred2a [Erdős and Dosztányi, 2020, Mészáros et al., 2018] using default options *long disorder* for entire proteins. We calculated the average disorder score of the whole sequence and percentage of residues predicted to be disordered. The percentage of disorder was calculated by taking the amount of disordered residues (disorder score > 0.5) and dividing it by the sequence length of the protein. We also predicted average disorder and percentage of disordered residues with a disorder threshold of 0.8 (**Figure S1**). A python script was used to automate predictions and disorder proportion for all candidates. We performed α-helix and β-sheet predictions to verify the amount of disordered residues predicted by IUPred2a. Secondary structure predictions were performed with Porter 5.0 (SS3) [Torrisi et al., 2018, 2019]. The predicted secondary structure elements for each residue were counted with a Javascript and divided by the total number of residues to obtain a percentage score for each structural element. DnaK binding sites were predicted using the ChaperISM suite (v1) in quantitative mode with default settings [Gutierres et al., 2020].

### Experimental Methods

#### Cloning of putative *de novo* candidates

Putative *de novo* candidates were synthesised as strings DNA from Twist Bioscience, San Francisco, codon optimzed for *E.coli* and without restriction sites used for cloning (BamH1, HindIII, NcoI, XhoI) inside the sequence. The wild-type DNA for Atlas was provided by Geoff Findlay. To introduce restriction sites at the ends we used different primers (a fasta file containing the DNA sequences and primer used can be found online on Zenodo (https://doi.org/10.5281/zenodo.6283284). For cloning into pHAT2 vector (N-terminal 6xHis) we used restriction enzymes combination of BamHI/XhoI + HindIII, for pETM-30 (N-terminal 6xHis-GST-TEV) we used NcoI+HindIII. Both vectors were from the EMBL vector database, Heidelberg, introduced stop-codon was TAA for all constructs. We digested the PCR product with both restriction enzymes respectively (FastDigest, Thermo Scientific) for 3 h at 37 C. Digest of the vector (1 h, 37 C) was purified from agarose gel (Zymo Research). We ligated both with an insert:vector ratio of 1:4 using Ligase (Thermo Scientific; 1 h, 22 C). The ligation mix was purified (Zymo Research) and 2 μL of the purified reaction mix was used to transform into 50 μL of chemically competent *E. coli* TOP10 cells. Cells were incubated for 30 min on ice, followed by a 90 sec heat-shock at 42 C. 500 μL of LB-Media (5 g yeast extract, 6 g tryptone, 5 g NaCl) was added for recovery and incubated for 1 h at 37 C. After incubation the resuspended cell pellets were plated on LB-agar containing 50 μg/mL ampicillin (AMP, Carl Roth, pHAT2, EMBL vector database) or Kanamycin (KAN, Carl Roth, pETM-30, EMBL vector database) and incubated at 37 C over night.

Successful transformation was verified by colony PCR and sequencing at Microsynth, Seqlab, Germany. The plasmid DNA bearing the chaperone combination GroEL/ES (pGro7) or DnaK/J/GrpE (pKJE) from Takara Biotech chaperone kit [Nishihara et al., 1998, 2000] were first transformed into *E coli* Top10 cells and then into expression strains (BL21 Star™ (DE3) and T7 Express). Chaperone plasmid bearing cells were made chemically competent (Inoue-method) [Inoue et al., 1990, Sambrook and Russell, 2006] and used for transformation with the plasmid containing the target protein sequence. Final expression cells contained two plasmids: chaperone plasmid and target protein plasmid. The chaperone plasmids are chloramphenicol (CAM) resistant, so the double plasmid cells are either AMP+CAM (pHAT2, N-terminal 6xHistag) or KAN+CAM (pETM-30, N-terminal GST-tag) resistant.

#### Test-Expression of candidate *de novo* proteins

To identify in which strain and plasmid proteins were expressed we performed test expressions. 10 mL of LB+AMP+CAM or LB+KAN+CAM were inoculated from a glycerol stock of all three expression cells bearing both plasmids (target protein and chaperone) and grown until turbid (6-8 h, 37). We split the solutions into 3×3 mL and incubated for 30 min at different temperatures (37 C, 28 C, and 20 C) before adding IPTG (Carl Roth) for a final concentration of 0.5 mM and shaking over night. When using the cells with chaperone plasmids we made the following adjustment: L-arabinose (final concentration 3 mM, Carl Roth) was added from the beginning for immediate induction of chaperone expression. Therefore, after inducing the *de novo* protein expression with IPTG the chaperones were already present in order to help folding the *de novo* proteins.

500 of each cell culture were centrifuged (15000 rpm, 2min). Pellets were re-suspended and lysed in 50of a mix of Bugbuster and Lysonase (both Merck AG) through vortexing for 10 min. After centrifugation the supernatant was mixed with the same volume of SDS-loading buffer (standard). The pellet was resuspended in 5x diluted Bugbuster, centrifuged and resuspended in 50 SDS-loading buffer. 15 of each fraction was loaded on an SDS-PAGE, either 10 % Bis-Tris or 12.5% TGS, run on 200V for 50min and dyed using ReadyBlueTM staining.

For the final western blots the determined optimal combination of strain, expression vector and chaperone plasmid was used. 20 mL cultures of 2YT+AMP+CAM or 2YT+KAN+CAM were inoculated with 1mL of the overnight culture. L-arabinose (final concentration 3 mM) was added to the samples, but not to the control without chaperones and grown at 37C, 180 rpm for 4-6 hours until turbid. The cultures were incubated at 28 C, 180 rpm for 30 min before induction with IPTG (final concentration 0.5 mM) and incubated overnight under these conditions. Final samples were harvested and handled as prior performed test-expressions.

#### Western blot

The SDS-PAGEs were run as described above but without ReadyBlueTMstaining. The gel was equilibrated in transfer buffer (20 % Methanol) for a few seconds. A polyvinylidene fluoride (PVDF) membrane with a pore size of 0.22 m was activated by methanol (2 min) and equilibrated in transfer buffer.The semi-dry transfer was performed at 25V for 30 minutes using the BioRad standard protocol. The membrane was blocked at room temperature for 1 hour using 5 % bovine serum albumin BSA in phosphate-buffered saline with tween (PBS-T) then washed in PBS-T and incubated for 1 hour with anti-His antibody (MA1-21315-HRP) diluted 1:500. For chemiluminescence 0.5 mL luminol was mixed with 0.5 mL peroxide and distributed evenly on the membrane.

#### Mass spectrometry

Tryptic digest followed by mass spectrometry for peptide detection of the candidate proteins was performed by the Core Unit Proteomics group of Prof. Dr. Simone König, UKM Muenster.

## Acknowledgements

A.L., E.BB and M.A. were funded by the Volkswagen Stiftung (VWF), grant code 98183. K.B. was funded by the Deutsche Forschungsgemeinschaft (DFG, German Research Foundation) — 281125614/GRK2220. We thank Prof. Geoff Findlay (College of the Holy Cross, Boston, Massachusetts) for the Atlas WT-sequence, Anne Diehl (FMP Berlin) for the BL21 Star™ (DE3) and T7 Express cells, our Master students Kai Köstler and Roman Schauerte for their help in the lab, and Prof. Simone König from the Core Unit Proteomic facility for performing the tryptic digest and mass spectrometry. We thank Mark Harrison for comments on the manuscript.

## Author Contributions

A.L. + E.BB. designed research; M.A., L.A.E., and A.L. performed cloning and expression; M.A. and L.A.E. performed western blots. M.A., K.B. and L.A.E. performed predictions. All authors wrote and approved the final version of the manuscript.

## References

Diethard Tautz and Tomislav Domazet-Lošo. The Evolutionary Origin of Orphan Genes. Nature Reviews Genetics, 12(10):692–702, October 2011. ISSN 1471-0056. doi: 10.1038/nrg3053.

Aoife McLysaght and Laurence D. Hurst. Open Questions in the Study of de Novo Genes: What, How and Why. Nature Reviews Genetics, 17(9):567–578, September 2016. ISSN 1471-0064. doi: 10.1038/nrg.2016.78.

Jonathan F Schmitz and Erich Bornberg-Bauer. Fact or Fiction: Updates on How Protein-Coding Genes Might Emerge de Novo from Previously Non-Coding DNA. F1000Research, 6:57, January 2017. ISSN 2046-1402. doi: 10.12688/f1000research.10079.1.

Stephen Branden Van Van Oss and Anne-Ruxandra Carvunis. De Novo Gene Birth. PLOS Genetics, 15(5):e1008160, May 2019. ISSN 1553-7404. doi: 10.1371/journal.pgen.1008160.

Christian Rödelsperger, Neel Prabh, and Ralf J. Sommer. New Gene Origin and Deep Taxon Phylogenomics: Opportunities and Challenges. Trends in genetics: TIG, 35(12):914–922, December 2019. ISSN 0168-9525. doi: 10.1016/j.tig.2019.08.007.

Erich Bornberg-Bauer, Klara Hlouchova, and Andreas Lange. Structure and function of naturally evolved de novo proteins. Current Opinion in Structural Biology, 68:175–183, June 2021. ISSN 1879-033X. doi: 10.1016/j.sbi.2020.11.010.

Brennen Heames, Filip Buchel, Margaux Aubel, Vyacheslav Tretyachenko, Andreas Lange, Erich Bornberg-Bauer, and Klara Hlouchova. Experimental characterisation of de novo proteins and their unevolved random-sequence counterparts. bioRxiv, 2022. doi: 10.1101/2022.01.14.476368.

David A. Liberles, Grigory Kolesov, and Katharina Dittmar. Understanding Gene Duplication Through Biochemistry and Population Genetics. In Evolution after Gene Duplication, pages 1–21. John Wiley & Sons, Ltd, 2011. ISBN 978-0-470-61990-2. doi: 10.1002/9780470619902.ch1.

Susumu Ohno. Evolution by Gene Duplication. Springer-Verlag, 1970. ISBN 9780387052250. doi: 10.1002/tera.1420090224.

Erich Bornberg-Bauer and M. Mar Albà. Dynamics and Adaptive Benefits of Modular Protein Evolution. Current Opinion in Structural Biology, 23(3):459–466, June 2013. ISSN 1879-033X. doi: 10.1016/j.sbi.2013.02.012.

Nikolaos Vakirlis, Anne-Ruxandra Carvunis, and Aoife McLysaght. Synteny-based analyses indicate that sequence divergence is not the main source of orphan genes. eLife, 9: e53500, feb 2020. ISSN 2050-084X. doi: 10.7554/eLife.53500.

David J. Begun, Heather A. Lindfors, Melissa E. Thompson, and Alisha K. Holloway. Recently Evolved Genes Identified From Drosophila Yakuba and D. Erecta Accessory Gland Expressed Sequence Tags. Genetics, 172(3):1675–1681, March 2006. ISSN 0016-6731, 1943-2631. doi: 10.1534/genetics.105.050336.

Jing Cai, Ruoping Zhao, Huifeng Jiang, and Wen Wang. De Novo Origination of a New Protein-Coding Gene in Saccharomyces Cerevisiae. Genetics, 179(1):487–496, May 2008. ISSN 0016-6731, 1943-2631. doi: 10.1534/genetics.107.084491.

Rafik Neme and Diethard Tautz. Phylogenetic Patterns of Emergence of New Genes Support a Model of Frequent de Novo Evolution. BMC Genomics, 14(1):117, February 2013. ISSN 1471-2164. doi: 10.1186/1471-2164-14-117.

Aoife McLysaght and Daniele Guerzoni. New Genes from Non-Coding Sequence: The Role of de Novo Protein-Coding Genes in Eukaryotic Evolutionary Innovation. Philosophical Transactions of the Royal Society B: Biological Sciences, 370(1678):20140332, September 2015. ISSN 0962-8436, 1471-2970. doi: 10.1098/rstb.2014.0332.

Christian Schlötterer. Genes from Scratch – the Evolutionary Fate of de Novo Genes. Trends in Genetics, 31(4):215–219, April 2015. ISSN 0168-9525. doi: 10.1016/j.tig.2015.02.007.

Jonathan F. Schmitz, Kristian K. Ullrich, and Erich Bornberg-Bauer. Incipient de Novo Genes Can Evolve from Frozen Accidents That Escaped Rapid Transcript Turnover. Nature Ecology & Evolution, 2(10):1626–1632, October 2018. ISSN 2397-334X. doi: 10.1038/s41559-018-0639-7.

Nikolaos Vakirlis, Alex S. Hebert, Dana A. Opulente, Guillaume Achaz, Chris Todd Hittinger, Gilles Fischer, Joshua J. Coon, and Ingrid Lafontaine. A Molecular Portrait of De Novo Genes in Yeasts. Molecular Biology and Evolution, 35(3):631–645, March 2018. ISSN 0737-4038. doi: 10.1093/molbev/msx315.

Neel Prabh and Christian Rödelsperger. De Novo, Divergence, and Mixed Origin Contribute to the Emergence of Orphan Genes in Pristionchus Nematodes. G3: Genes, Genomes, Genetics, page g3.400326.2019, May 2019. ISSN 2160-1836. doi: 10.1534/g3.119.400326.

Li Zhang, Yan Ren, Tao Yang, Guangwei Li, Jianhai Chen, Andrea R. Gschwend, Yeisoo Yu, Guixue Hou, Jin Zi, Ruo Zhou, Bo Wen, Jianwei Zhang, Kapeel Chougule, Muhua Wang, Dario Copetti, Zhiyu Peng, Chengjun Zhang, Yong Zhang, Yidan Ouyang, Rod A. Wing, Siqi Liu, and Manyuan Long. Rapid Evolution of Protein Diversity by de Novo Origination in Oryza. Nature Ecology & Evolution, 3(4):679, April 2019. ISSN 2397-334X. doi: 10.1038/s41559-019-0822-5.

Brennen Heames, Jonathan Schmitz, and Erich Bornberg-Bauer. A Continuum of Evolving de Novo Genes Drives Protein-Coding Novelty in Drosophila. Journal of Molecular Evolution, 2020. doi: 10.1007/s00239-020-09939-z.

Daniel Dowling, Jonathan F. Schmitz, and Erich Bornberg-Bauer. Stochastic gain and loss of novel transcribed open reading frames in the human lineage. Genome Biology and Evolution, 12:2183 – 2195, 2020. doi: 10.1093/gbe/evaa194.

Anna Grandchamp, Katrin Berk, Elias Dohmen, and Erich Bornberg-Bauer. New genomic signals underlying the emergence of human proto-genes. bioRxiv, 2022. doi: 10.1101/2022.01.04.474757.

Tomislav Domazet-Lošo, Anne-Ruxandra Carvunis, M. Mar Albà, Martin Sebastijan Šestak, Robert Bakarić, Rafik Neme, and Diethard Tautz. No Evidence for Phylostratigraphic Bias Impacting Inferences on Patterns of Gene Emergence and Evolution. Molecular Biology and Evolution, 34(4):843–856, April 2017. ISSN 0737-4038. doi: 10.1093/molbev/msw284.

Andreas Lange, Prajal H. Patel, Brennen Heames, Adam M. Damry, Thorsten Saenger, Colin J. Jackson, Geoffrey D. Findlay, and Erich Bornberg-Bauer. Structural and functional characterization of a putative de novo gene in Drosophila. Nature Communications, 12(1): 1667, March 2021. ISSN 2041-1723. doi: 10.1038/s41467-021-21667-6.

Dixie Bungard, Jacob S. Copple, Jing Yan, Jimmy J. Chhun, Vlad K. Kumirov, Scott G. Foy, Joanna Masel, Vicki H. Wysocki, and Matthew H. J. Cordes. Foldability of a Natural De Novo Evolved Protein. Structure, 25(11):1687–1696.e4, November 2017. ISSN 0969-2126. doi: 10.1016/j.str.2017.09.006.

Nobuhiko Tokuriki and Dan S. Tawfik. Chaperonin overexpression promotes genetic variation and enzyme evolution. Nature, 459:668–673, 2009a. doi: 10.1038/nature08009.

Nobuhiko Tokuriki and Dan S. Tawfik. Protein dynamism and evolvability. Science, 324:203 – 207, 2009b. doi: 10.1126/science.1169375.

Nobuhiko Tokuriki and Dan S. Tawfik. Stability effects of mutations and protein evolvability. Current opinion in structural biology, 19 5:596–604, 2009c. doi: 10.1016/j.sbi.2009.08.003.

Colin Jackson, Agnes Toth-Petroczy, Rachel Kolodny, Florian Hollfelder, Monika Fuxreiter, Shina Caroline Lynn Kamerlin, and Nobuhiko Tokuriki. Adventures on the routes of protein evolution — in memoriam dan salah tawfik (1955 - 2021). Journal of Molecular Biology, page 167462, 2022. ISSN 0022-2836. doi: 10.1016/j.jmb.2022.167462.

Misha Soskine and Dan S. Tawfik. Mutational effects and the evolution of new protein functions. Nature Reviews Genetics, 11:572–582, 2010. doi: 10.1038/nrg2808.

Vyacheslav Tretyachenko, Jiří Vymětal, Lucie Bednárová, Vladimír Kopecký, Kateřina Hofbauerová, Helena Jindrová, Martin Hubálek, Radko Souček, Jan Konvalinka, Jiří Von-drášek, and Klára Hlouchová. Random Protein Sequences Can Form Defined Secondary Structures and Are Well-Tolerated in Vivo. Scientific Reports, 7(1):15449, November 2017. ISSN 2045-2322. doi: 10.1038/s41598-017-15635-8.

Brigitte Gasser, Markku Saloheimo, Ursula Rinas, Martin Dragosits, Escarlata Rodríguez-Carmona, Kristin Baumann, Maria Giuliani, Ermenegilda Parrilli, Paola Branduardi, Christine Lang, Danilo Porro, Pau Ferrer, Maria Luisa Tutino, Diethard Mattanovich, and Antonio Villaverde. Protein folding and conformational stress in microbial cells producing recombinant proteins: a host comparative overview. Microbial Cell Factories, 7:11 – 11, 2008. doi: 10.1186/1475-2859-7-11.

Erich Bornberg-Bauer, Jonathan F. Schmitz, and Magdalena Heberlein. Emergence of de novo proteins from ‘dark genomic matter’ by ‘grow slow and moult’. Biochemical Society transactions, 43 5:867–73, 2015. doi: 10.1042/BST20150089.

Andrija Finka, Rayees U. H. Mattoo, and Pierre Goloubinoff. Experimental milestones in the discovery of molecular chaperones as polypeptide unfolding enzymes. Annual review of biochemistry, 85:715–42, 2016. doi: 10.1146/annurev-biochem-060815-014124.

David S. Libich, Vitali Tugarinov, and G. Marius Clore. Intrinsic unfoldase/foldase activity of the chaperonin groel directly demonstrated using multinuclear relaxation-based nmr. Proceedings of the National Academy of Sciences, 112:8817 – 8823, 2015. doi: 10.1073/pnas.1510083112.

Zong Lin, Damian Madan, and Hays S. Rye. Groel stimulates protein folding through forced unfolding. Nature Structural &Molecular Biology, 15:303–311, 2008. doi: 10.1038/nsmb.1394.

Jeffrey G. Thomas, Amanda Ayling, and Francois. Baneyx. Molecular chaperones, folding catalysts, and the recovery of active recombinant proteins from E. coli. To fold or to refold. Appl Biochem Biotechnol, 66(3):197–238, Jun 1997. doi: 10.1007/BF02785589.

Hartwig Schröder, Thomas Langer, F. Ulrich Hartl, and Bernd Bukau. Dnak, dnaj and grpe form a cellular chaperone machinery capable of repairing heat-induced protein damage. The EMBO Journal, 12, 1993. doi: 10.1002/j.1460-2075.1993.tb06097.x.

Sandeep Savitaprakash Sharma, Paolo De Los Rios, Philipp Christen, Ariel Lustig, and Pierre Goloubinoff. The kinetic parameters and energy cost of the hsp70 chaperone as a polypeptide unfoldase. Nature chemical biology, 6 12:914–20, 2010. doi: 10.1038/nchembio.455.

Yujin E. Kim, Mark S. Hipp, Andreas Bracher, Manajit Hayer-Hartl, and F. Ulrich Hartl. Molecular chaperone functions in protein folding and proteostasis. Annual review of biochemistry, 82:323–55, 2013. doi: 10.1146/annurev-biochem-060208-092442.

Alireza Mashaghi, Sergey Bezrukavnikov, David P. Minde, Anne S Wentink, Roman Kityk, Beate Zachmann-Brand, Matthias P. Mayer, Günter Kramer, Bernd Bukau, and Sander J. Tans. Alternative modes of client binding enable functional plasticity of hsp70. Nature, 539:448–451, 2016. doi: 10.1038/nature20137.

Pierre Goloubinoff, Anthony A. Gatenby, and George H. Lorimer. Groe heat-shock proteins promote assembly of foreign prokaryotic ribulose bisphosphate carboxylase oligomers in escherichia coli. Nature, 337:44–47, 1989. doi: 10.1038/337044a0.

Emily L. Rivard, Andrew G. Ludwig, Prajal H. Patel, Anna Grandchamp, Sarah E. Arnold, Alina Berger, Emilie M. Scott, Brendan J. Kelly, Grace C. Mascha, Erich Bornberg-Bauer, and Geoffrey D. Findlay. A putative de novo evolved gene required for spermatid chromatin condensation in Drosophila melanogaster. PLOS Genetics, 17(9):e1009787, September 2021. ISSN 1553-7404. doi: 10.1371/journal.pgen.1009787.

Gábor Erdős and Zsuzsanna Dosztányi. Analyzing Protein Disorder with IUPred2A. Current Protocols in Bioinformatics, 70(1):e99, 2020. ISSN 1934-340X. doi: 10.1002/cpbi.99.

Bálint Mészáros, Gábor Erdős, and Zsuzsanna Dosztányi. IUPred2A: Context-dependent prediction of protein disorder as a function of redox state and protein binding. Nucleic Acids Research, 46(W1):W329–W337, July 2018. ISSN 0305-1048. doi: 10.1093/nar/gky384.

Mirko Torrisi, Manaz Kaleel, and Gianluca Pollastri. Porter 5: fast, state-of-the-art ab initio prediction of protein secondary structure in 3 and 8 classes. bioRxiv, 2018. doi: 10.1101/289033.

Mirko Torrisi, Manaz Kaleel, and Gianluca Pollastri. Deeper profiles and cascaded recurrent and convolutional neural networks for state-of-the-art protein secondary structure prediction. Scientific Reports, 9, 2019. doi: 10.1038/s41598-019-48786-x.

Bryan A. Moyers and Jianzhi Zhang. Phylostratigraphic Bias Creates Spurious Patterns of Genome Evolution. Molecular Biology and Evolution, 32(1):258–267, January 2015. ISSN 0737-4038, 1537-1719. doi: 10.1093/molbev/msu286.

Caroline M. Weisman, Andrew W. Murray, and Sean R. Eddy. Mixing genome annotation methods in a comparative analysis inflates the apparent number of lineage-specific genes. bioRxiv, 2022. doi: 10.1101/2022.01.13.476251.

Fakhri Saïda, Marc Uzan, Benoît Odaert, and Francois Bontems. Expression of highly toxic genes in E. coli: Special strategies and genetic tools. Current Protein & Peptide Science, 7(1):47–56, February 2006. ISSN 1389-2037. doi: 10.2174/138920306775474095.

Fakhri Saïda. Overview on the expression of toxic gene products in Escherichia coli. Current Protocols in Protein Science, Chapter 5:Unit 5.19, November 2007. ISSN 1934-3663. doi: 10.1002/0471140864.ps0519s50.

Germán L. Rosano and Eduardo A. Ceccarelli. Recombinant protein expression in Escherichia coli: Advances and challenges. Frontiers in Microbiology, 5:172, 2014. ISSN 1664-302X. doi: 10.3389/fmicb.2014.00172.

New England Biolabs. Datasheet for t7 express competent e. coli (high efficiency) (c2566; lot 18). https://www.nebiolabs.com.au/-/media/catalog/datacards-or-manuals/c2566datasheet-lot18.pdf?rev=234841213ece47a48f9da8de895ca3db&hash=CB482DAE0DA6659F3B5B7618615B4902 (Accessed on 02/24/2022).

Sandra Harper and David W. Speicher. Purification of proteins fused to glutathione Stransferase. Methods in Molecular Biology (Clifton, N.J.), 681:259–280, 2011. ISSN 1940-6029. doi: 10.1007/978-1-60761-913-0_14.

Tatsuya Niwa, Takashi Kanamori, Takuya Ueda, and Hideki Taguchi. Global analysis of chaperone effects using a reconstituted cell-free translation system. Proc Natl Acad Sci U S A, 109(23):8937–8942, Jun 2012. doi: 10.1073/pnas.1201380109.

Gustav N. Sundell and Ylva Ivarsson. Interaction analysis through proteomic phage display. BioMed Research International, 2014:176172, 2014. ISSN 2314-6141. doi: 10.1155/2014/176172.

Ylva Ivarsson, Roland Arnold, Megan McLaughlin, Satra Nim, Rakesh Joshi, Debashish Ray, Bernard Liu, Joan Teyra, Tony Pawson, Jason Moffat, Shawn Shun-Cheng Li, Sachdev S. Sidhu, and Philip M. Kim. Large-scale interaction profiling of PDZ domains through proteomic peptide-phage display using human and viral phage peptidomes. Proceedings of the National Academy of Sciences, 111(7):2542–2547, February 2014. ISSN 0027-8424, 1091-6490. doi: 10.1073/pnas.1312296111.

Muhammad Ali, Leandro Simonetti, and Ylva Ivarsson. Screening Intrinsically Disordered Regions for Short Linear Binding Motifs. In Birthe B. Kragelund and Karen Skriver, editors, Intrinsically Disordered Proteins: Methods and Protocols, Methods in Molecular Biology, pages 529–552. Springer US, New York, NY, 2020. ISBN 978-1-07-160524-0. doi: 10.1007/978-1-0716-0524-0_27.

Yi-Hsiang Chiu, K. T. Ko, Tzu-Jing Yang, Kuen-Phon Wu, Meng-Ru Ho, Piotr Draczkowski, and Shang-Te Danny Hsu. Direct visualization of a 26 kda protein by cryo-electron microscopy aided by a small scaffold protein. Biochemistry, 2021. doi: 10.1021/acs.biochem.0c00961.

John M. Jumper, Richard Evans, Alexander Pritzel, Tim Green, Michael Figurnov, Olaf Ronneberger, Kathryn Tunyasuvunakool, Russ Bates, Augustin Zídek, Anna Potapenko, Alex Bridgland, Clemens Meyer, Simon A A Kohl, Andy Ballard, Andrew Cowie, Bernardino Romera-Paredes, Stanislav Nikolov, Rishub Jain, Jonas Adler, Trevor Back, Stig Petersen, David A. Reiman, Ellen Clancy, Michal Zielinski, Martin Steinegger, Michalina Pacholska, Tamas Berghammer, Sebastian Bodenstein, David Silver, Oriol Vinyals, Andrew W. Senior, Koray Kavukcuoglu, Pushmeet Kohli, and Demis Hassabis. Highly accurate protein structure prediction with alphafold. Nature, 596:583 – 589, 2021. doi: 10.1038/s41586-021-03819-2.

Minkyung Baek, Frank DiMaio, Ivan Anishchenko, Justas Dauparas, Sergey Ovchinnikov, Gyu Rie Lee, Jue Wang, Qian Cong, Lisa N Kinch, R Dustin Schaeffer, et al. Accurate prediction of protein structures and interactions using a three-track neural network. Science, 373(6557):871–876, 2021. doi: 10.1126/science.abj8754.

Kiersten M Ruff and Rohit V Pappu. Alphafold and implications for intrinsically disordered proteins. Journal of Molecular Biology, 433(20):167208, 2021. doi: 10.1016/j.jmb.2021.167208.

Vivian Monzon, Daniel H Haft, and Alex Bateman. Folding the unfoldable: using alphafold to explore spurious proteins. Bioinformatics Advances, 2(1):vbab043, 2022. doi: 10.1093/bioadv/vbab043.

M. B. B. Gutierres, Cristina Bonorino, and Maurício Menegatti Rigo. Chaperism: improved chaperone binding prediction using position-independent scoring matrices. Bioinformatics, 2020. doi: 10.1093/bioinformatics/btz670.

Kazuyo Nishihara, Masaaki Kanemori, Masanari Kitagawa, Hideki Yanagi, and Takashi Yura. Chaperone coexpression plasmids: Differential and synergistic roles of dnak-dnaj-grpe and groel-groes in assisting folding of an allergen of japanese cedar pollen, cryj2, inescherichia coli. Applied and Environmental Microbiology, 64:1694 – 1699, 1998. doi: 10.1128/AEM.64.5.1694-1699.1998.

Kazuyo Nishihara, Masaaki Kanemori, Hideki Yanagi, and Takashi Yura. Overexpression of trigger factor prevents aggregation of recombinant proteins in escherichia coli. Applied and Environmental Microbiology, 66:884 – 889, 2000. doi: 10.1128/AEM.66.3.884-889.2000.

Hiroaki Inoue, Hiroshi Nojima, and Hiroto Okayama. High efficiency transformation of Escherichia coli with plasmids. Gene, 96(1):23–28, November 1990. ISSN 0378-1119. doi: 10.1016/0378-1119(90)90336-p.

Joseph Sambrook and David W. Russell. The Inoue Method for Preparation and Transformation of Competent E. Coli: “Ultra-Competent” Cells. Cold Spring Harbor Protocols, 2006 (1):pdb.prot3944, January 2006. ISSN 1940-3402, 1559-6095. doi: 10.1101/pdb.prot3944.

