## Supplementary figures and images for "Chaperones facilitate heterologous expression of naturally evolved putative *de novo* proteins"

### Figure S1

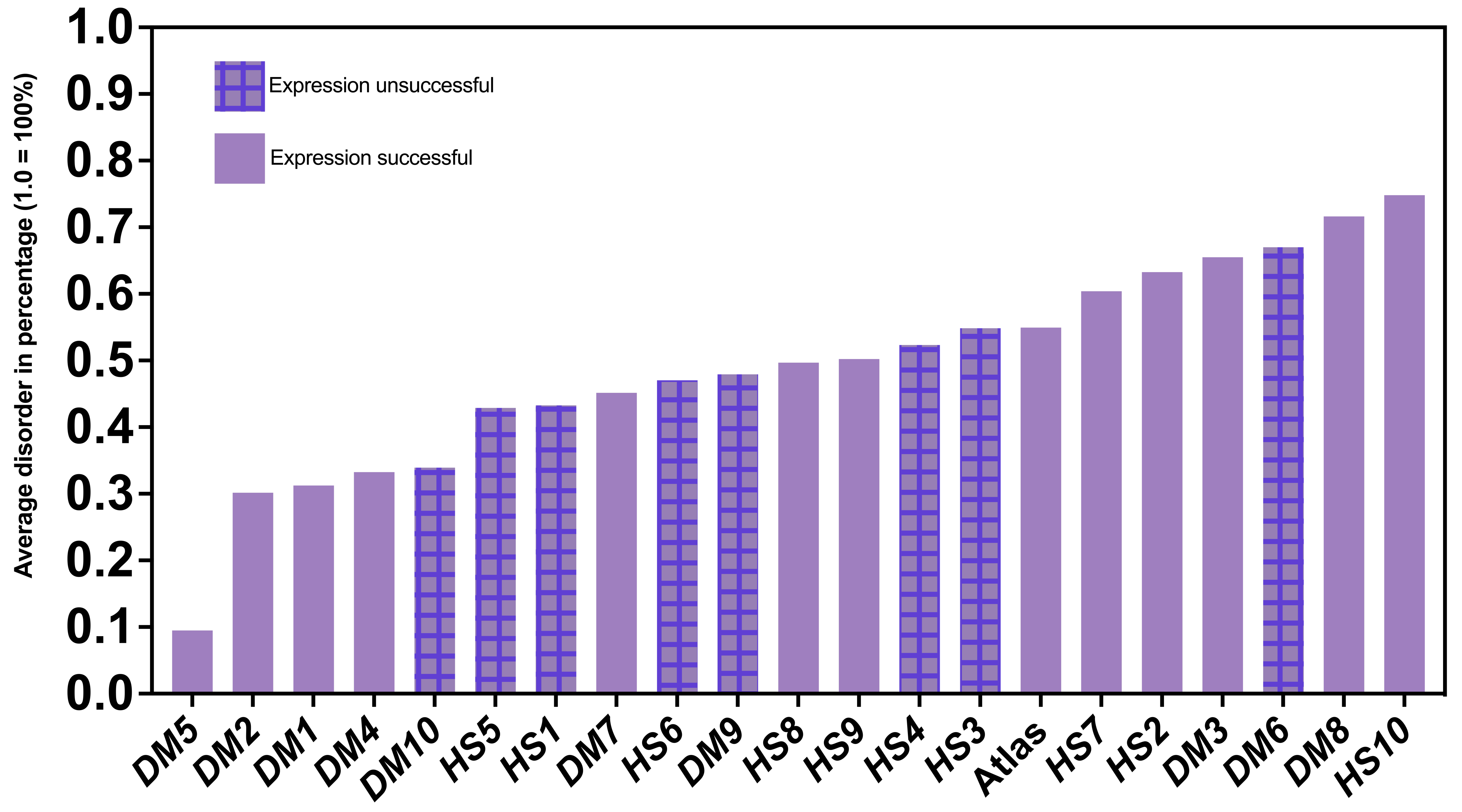

### Figure S2

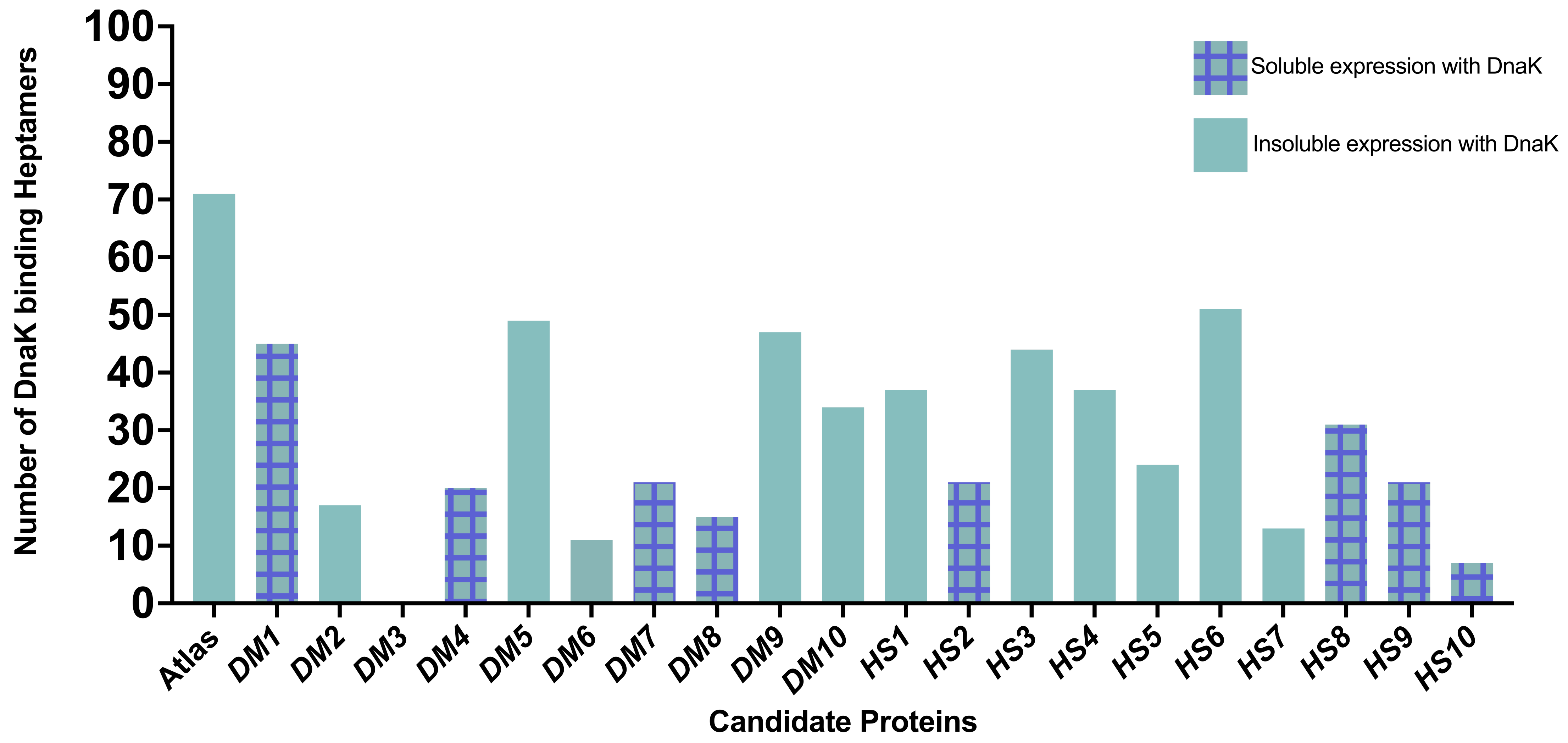

### Figure S3

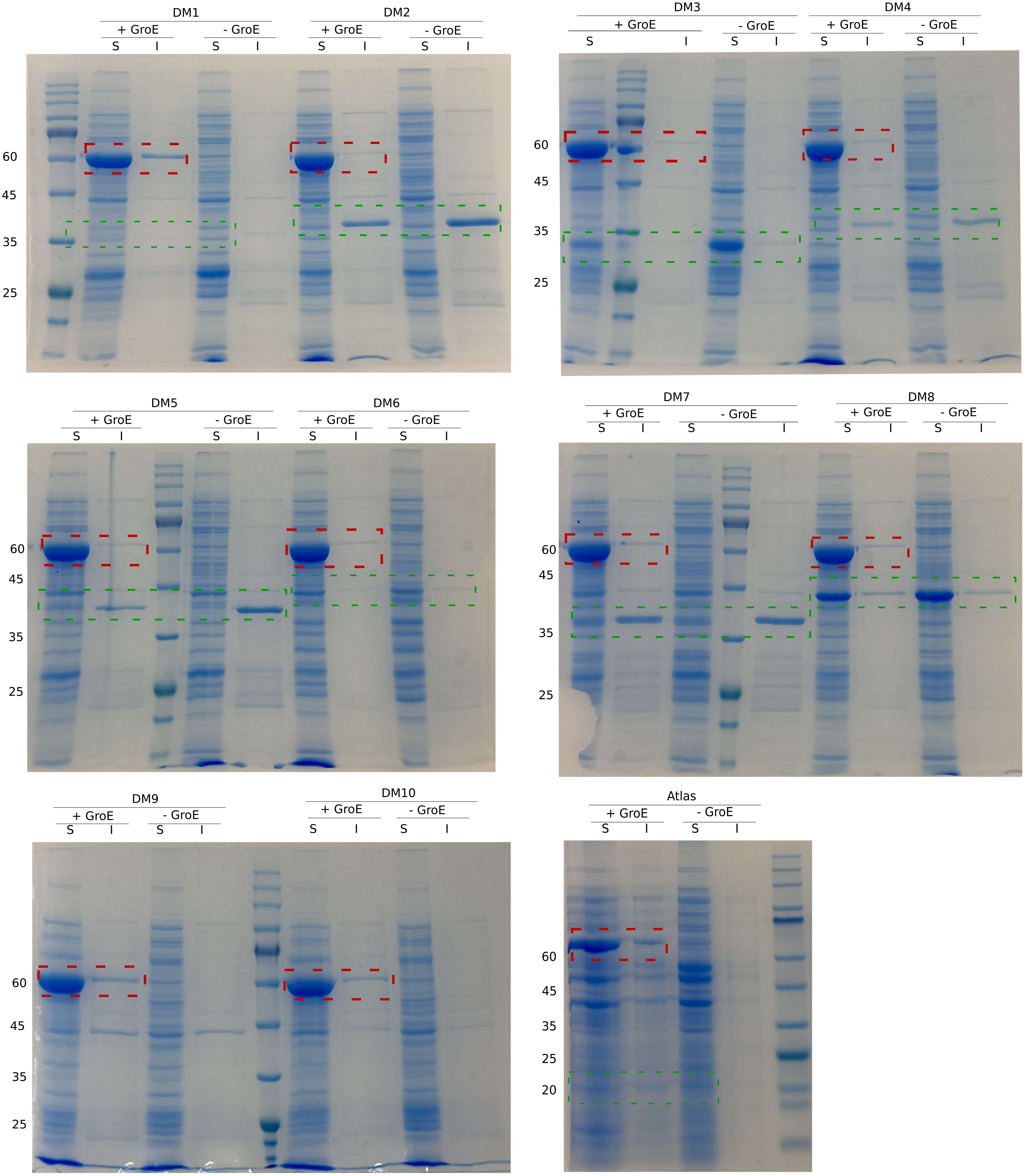

### Figure S4

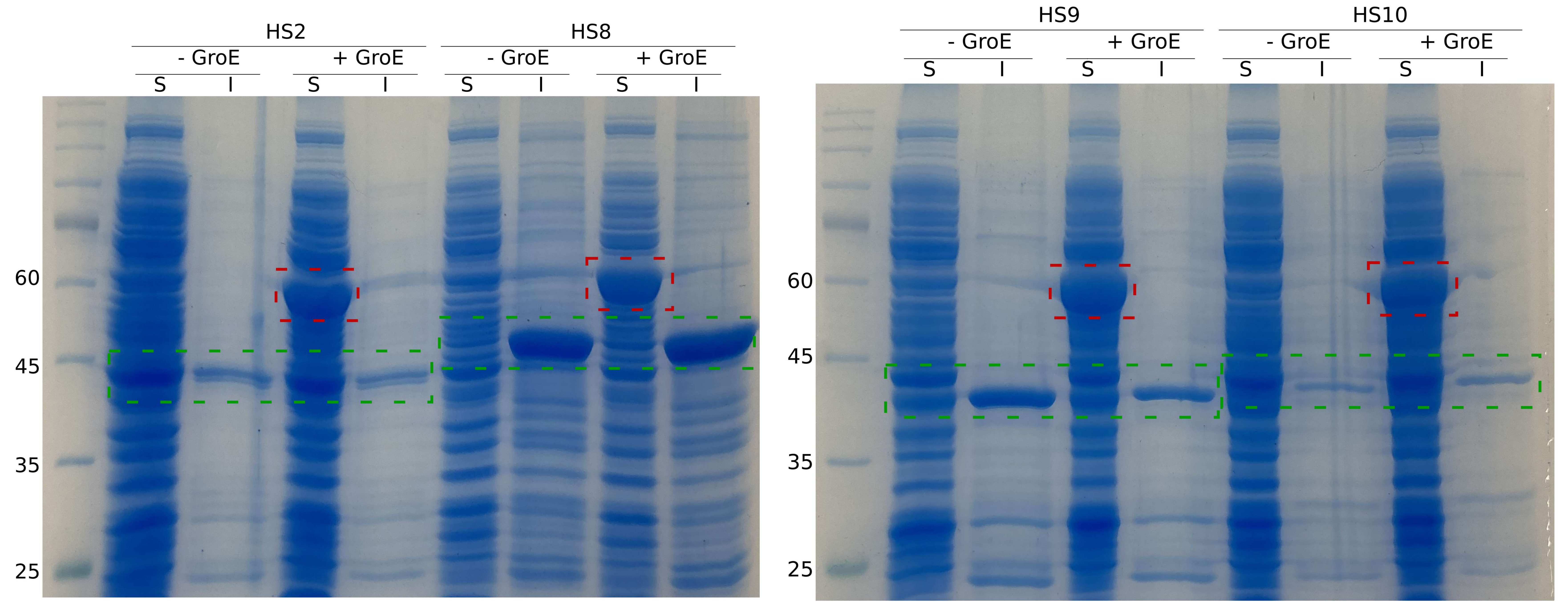
